# A data-driven redefinition of global biodiversity hotspots

**DOI:** 10.64898/2026.05.07.721789

**Authors:** Xin Liu, David Lindenmayer, Colin A. Chapman, Paul A. Garber, Rong Li, Cyril C. Grueter, Ruidong Wu, Yin Yang

## Abstract

The 36 global biodiversity hotspots harbor a disproportionate share of the world’s endemic species, making their conservation critical for planetary health. Traditionally hotspots were defined as ecoregions with ≥1,500 endemic vascular plant species and >70% natural habitat loss; this relied heavily on expert judgment, with subjective assessments of endemism and habitat loss applied. Challenges in defining endemism, quantifying habitat loss, and the global unevenness in available vascular plant data have hindered hotspot identification over the past decades. Here, we built a global dataset of 150,487 rare vascular plants, identified from 88.1% of the world’s known vascular species, and recognize hotspots based on their richness using three complementary conservation targets and algorithms. We then quantified natural habitat loss and habitat fragmentation using high-resolution remote sensing data and assessed the diversity and distribution of terrestrial vertebrates within these newly identified hotspots. Our data-driven method recovered all the 36 established global biodiversity hotspots, revised 17, and identified 11 new hotspots spanning diverse ecosystems across six continents. These 47 hotspots cover 26.63% of global land area, yet contain 83.8% of rare vascular plants, 92.4% of mammals, 96.1% of birds, 87.8% of reptiles, and 95.0% of amphibians. Collectively, they encompass >89% of terrestrial vertebrates classified as IUCN threatened species. Only 10 hotspots have undergone ≥70% habitat loss, and the lack of a consistent relationship with habitat fragmentation suggests that this criterion is not globally applicable. Effectively protecting ∼27% of Earth’s land could theoretically safeguard >89.8% of threatened terrestrial species and >67% of threatened terrestrial species hotspot area, assuming effective protection of the identified biodiversity hotspots. Targeted conservation efforts within these global biodiversity hotspots can meet the established biodiversity targets of the Kunming–Montreal Global Biodiversity Framework as well as post-2030 biodiversity targets. Most importantly, our framework enables conservation scientists to iteratively identify and update global biodiversity hotspots in step with growing global biodiversity data.

## Main

Biodiversity hotspots are globally recognized as priority conservation areas due to their unique vascular plant endemism, species richness, and high level of threat (*1*, *2*). The hotspot concept was first introduced by Norman Myers in 1988(*3*), and provides a strategic framework that emphasizes both biological uniqueness and the extent of habitat loss in conservation priority areas. The concept, as subsequently revised by Myers *et al*. in 2000 (*1*), identifies a region as a biodiversity hotspot if it contains at least 1,500 endemic vascular plant species and has lost over 70% of its original native vegetation. This hotspot framework has guided global conservation priorities for decades, leading to the identification of 36 terrestrial hotspots by 2016, including iconic biodiversity regions such as the Tropical Andes, Eastern Himalaya, Mediterranean Basin, Eastern Afromontane, and North American Coastal Plains (*1*, *4*). The hotspot concept has shaped conservation policy and investment, attracting more than US$300 million in past three decades from a single foundation alone, the Critical Ecosystem Partnership Fund (*5*).

### Constraints of Existing Hotspot Identification Criteria

Despite its broad adoption, the hotspot framework has notable limitations. Criteria for identifying local endemism rely heavily on subjective expert judgment and long-term accumulation of species inventories. Furthermore, definitions centered on habitat loss thresholds continue to lack scientific consensus regarding the quantitative delimitation of local endemism (*6*). Determining whether a species is truly endemic to a region is often complicated by incomplete distributional data, taxonomic uncertainties, and assessments of species ranges that are incompletely known or changing (*7*, *8*). In poorly surveyed or remote areas, limited inventories and misinterpretations of endemism (e.g., assuming endemic species are rare or narrowly distributed when they may be regionally common) can skew assessments of how well it represents biodiversity (*9*, *10*). Species richness also scales with area, making the spatial definition of endemism inconsistent (*10*). For example, in eastern Australian forests, endemic vascular plants are defined by having more than 50 occurrence records, with 95% restricted to two ecoregions, while in the North American Coastal Plains, endemics are defined by >90% of occurrences falling within the hotspot boundary (*4*). Therefore, a method for delineating endemic plant-rich regions that performs well in one hotspot may not be applicable to others, highlighting the need for globally consistent and data-driven criteria.

An additional problem is that the 70% natural habitat loss criterion used to define the degree of degradation is not only a very arbitrary criterion, but varies depending on data sources and evaluation methods. Different hotspot assessments use different and changing land cover datasets and thresholds, leading to inconsistent application of criteria across regions (*11*). Some regions of great ecological significance and which are renowned for their biodiversity, such as the Amazon (including the Guiana Highlands), the Congo basin, Papua New Guinea, and the Appalachian Mountains, have not been formally included as biodiversity hotspots because native vegetation loss has not reached 70% (*12–15*). Moreover, natural sparsely-vegetated systems such as grasslands, savannas, and deserts have frequently been misinterpreted in ecosystem ecology as degraded remnants or inherently species-poor habitats and have thus been overlooked as primary native biomes, despite their evolutionary distinctiveness and rich biodiversity (e.g., the Chihuahuan Desert and the dry-hot valley savanna systems of southwest China) (*16–18*). Thus, the current biodiversity hotspot framework overlooks areas of high conservation value, including some which may be close to a tipping point in terms of environmental change and impending extinctions (*19*, *20*).

### Toward Data-Driven Criteria for Hotspot Redefinition

Amid escalating global environmental challenges, such as climate change, species extinctions, habitat loss, and pollution (*2*, *21*, *22*), there is an urgent need to reassess and refine the concept of biodiversity hotspots to ensure that conservation strategies are scientifically robust and remain effective in guiding the use of limited conservation resources (*13*, *23*). Recent research has advocated for more objective, comprehensive, and data-driven approaches that integrate multiple biodiversity metrics, incorporate data at fine-scale spatial resolution, and account for dynamic habitat change (*23–25*).

Advances in biodiversity data collection and spatial analysis, including the digitization of herbarium records, remote sensing, and increasingly complete species distribution databases such as GBIF, eBird, and IUCN expert maps, now enable more accurate and scalable assessments of biodiversity patterns (*26*). These tools facilitate the identification of areas with high concentrations of rare or threatened species, providing opportunities to improve the precision and inclusivity of hotspot delineation.

In this study, we developed a global database of rare vascular plants (hereafter RVP) based on the rarity concept proposed by Enquist *et al* (*27*) who defines rare vascular plant species as those recorded from no more than five specimen collection localities (SPD ≤ 5) worldwide, or under a stricter criterion of no more than three (SPD ≤ 3) (*27*). Rarity reflects intrinsic species properties, such as restricted range, low occupancy rate, and/or small population size, providing scale-independent indicators suited to global prioritization (*27*, *28*). Rare species disproportionately support vulnerable ecosystem functions (*29*), yet can be sensitive to habitat loss, climate change, and demographic stochasticity, making their disappearance a more immediate signal of biodiversity decline than counts of endemic species (*30*, *31*). Moreover, a rarity-based framework mitigates endemism bias, whereby endemically-rich regions may be dominated by widespread or stable taxa, while globally rare species in less endemic-rich regions are overlooked (*32*). Thus, RVP can provide a more globally consistent and biologically meaningful basis for hotspot identification than endemism alone, and prioritizing rarity may improve both the fidelity of hotspot delineation and the effectiveness of global conservation strategies. Additionally, in 2011, Mittermeier *et al*. estimated that the 35 recognized biodiversity hotspots (excluding the North American Coastal Plains) collectively encompassed approximately 60% of all threatened mammal species, 63% of threatened bird species, and 79% of threatened amphibian species (*13*). Therefore, in our study, we reassessed how the newly identified biodiversity hotspots capture the overall richness and the number of threatened species across four major terrestrial vertebrate groups, mammals, birds, reptiles, and amphibians.

Specifically, we addressed three key questions: (1) Do rare vascular plants provide a novel and reliable basis for delineating biodiversity hotspots and capturing global biodiversity patterns; (2) Where are newly identified hotspots located, and do they satisfy the conventional criterion of ≥70% native habitat loss; and (3) How do newly revealed hotspots reshape conservation priorities for meeting future global biodiversity conservation targets. We also offer a robust and scalable approach for conservation scientists to iteratively identify and adjust global biodiversity hotspots as additional biodiversity data become available.

## Results

### Spatial Distribution Patterns of Rare Vascular Plants

The global distribution patterns of both RVP categories (SPD3 and SPD5) were similar, showing multi-center clustering, a bell-shaped latitudinal curve, and multiple peaks between 50°N and 50°S (Fig. 1). Low-latitude concentrations of RVP richness occur in Central America, Amazonia, the Andes and Cerrado of South America, the Guinean Forests and Congo Basin of Central Africa, the East African Mountains, eastern Madagascar, the Southeast Asian mainland, the Malay Archipelago, and New Guinea with its surrounding islands. We observed mid-latitude peaks of RVP richness in the Appalachian Mountains and eastern North America, Mediterranean Europe, the Qinling-Daba Mountains and southwestern mountains and valleys of China, the Himalayas, and Japan. As might be expected, the high-latitude regions harbor only limited RVP richness, notably the Alps and Caucasus in Europe and the Western Cape of South Africa.

**Fig. 1.**
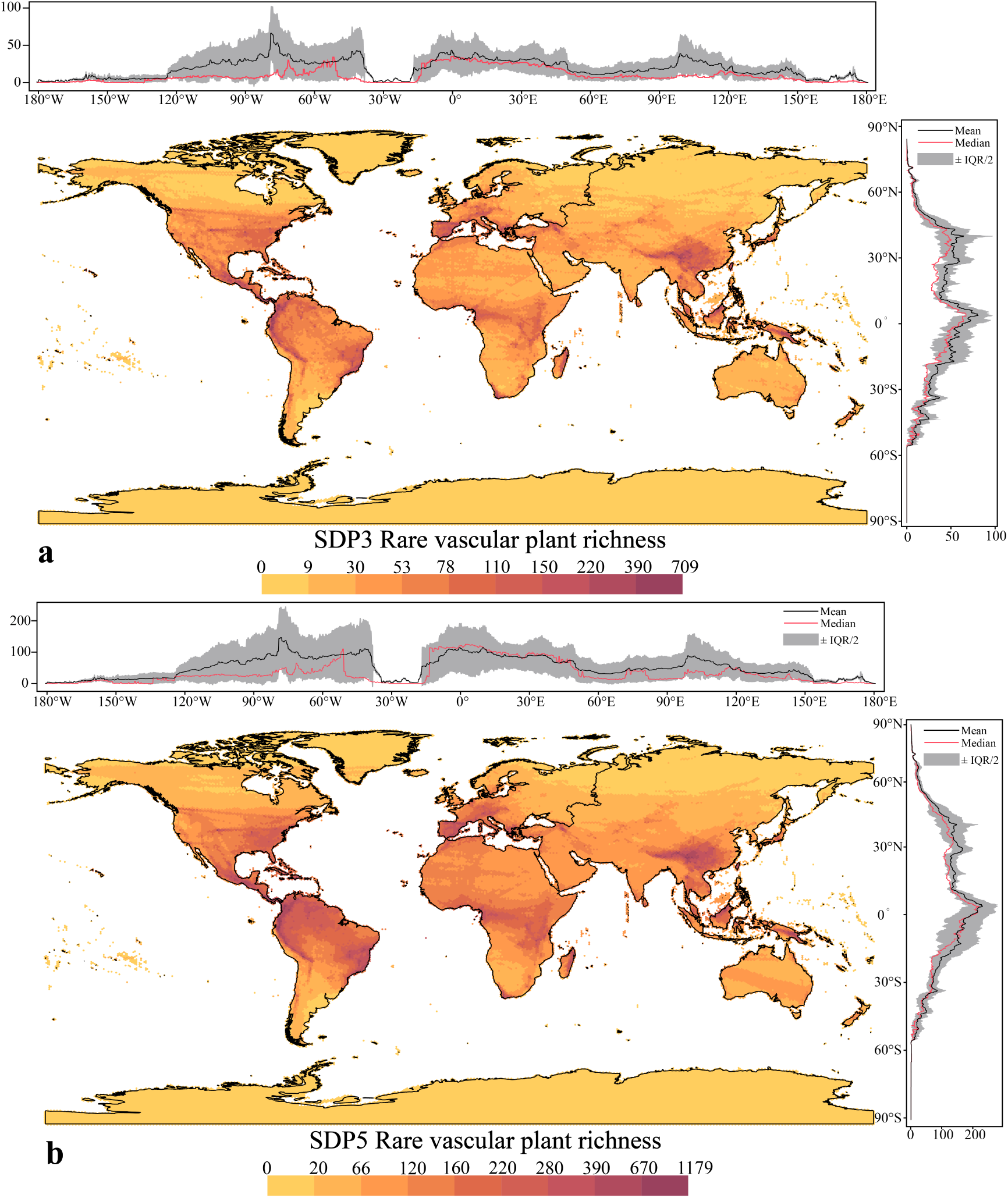
Global spatial distribution patterns of rare vascular plant species. (a) SPD3: Plant Species with Specimens Collected from ≤ Three Distribution Sites Worldwide. (b) SPD5: Plant Species with Specimens Collected from ≤ Five Distribution Sites Worldwide.

### Biodiversity Hotspots Delineated based on RVP

We identified 280 biodiversity hotspot ecoregions across WWF’s 867 terrestrial ecoregions by integrating RVP richness with distinct conservation objectives (details in Methods; Fig. 2a, Supplementary Data Fig. 1) (*33*). By integrating biome characteristics, topography, geomorphology, well-established biogeographical divisions, and climate zonation (Supplementary Document 1), we identified 35 global biodiversity hotspots (Fig. 2b). Among these hotspots, 23 overlapped with those demarcated those of Myers *et al*. (*1*). However, the land areas of three previously identified hotspots, the Southwest China Mountains, the Cerrado of South America, and the Eastern Afromontane, were substantially expanded (Supplementary Table 1), primarily through the integration of similar or adjacent biomes and associated topographic and geomorphological units from neighboring hotspot ecoregions. For example, we defined the Hengduan Mountains hotspot by integrating the Southeast Tibet Shrublands and Meadows and the Qilian Mountains Conifer Forests with the Southwest China Mountains hotspot from Myers *et al*.’s framework. This revised region encompasses the full range of ecosystems across the Hengduan Mountains. The remaining 12 global biodiversity hotspots are newly identified and span six continents (Fig. 2). Notably, the 35 identified biodiversity hotspots did not include biodiverse regions such as the eastern and western Australian forests and New Zealand. In contrast, the hotspots identified here using the Dobson algorithm^34^ aimed at protecting all RVP species encompass each of the 36 global biodiversity hotspots proposed under Myers *et al*.’ s framework (Extended Data Fig. 1 and Extended Data Fig. 4).

**Fig. 2.**
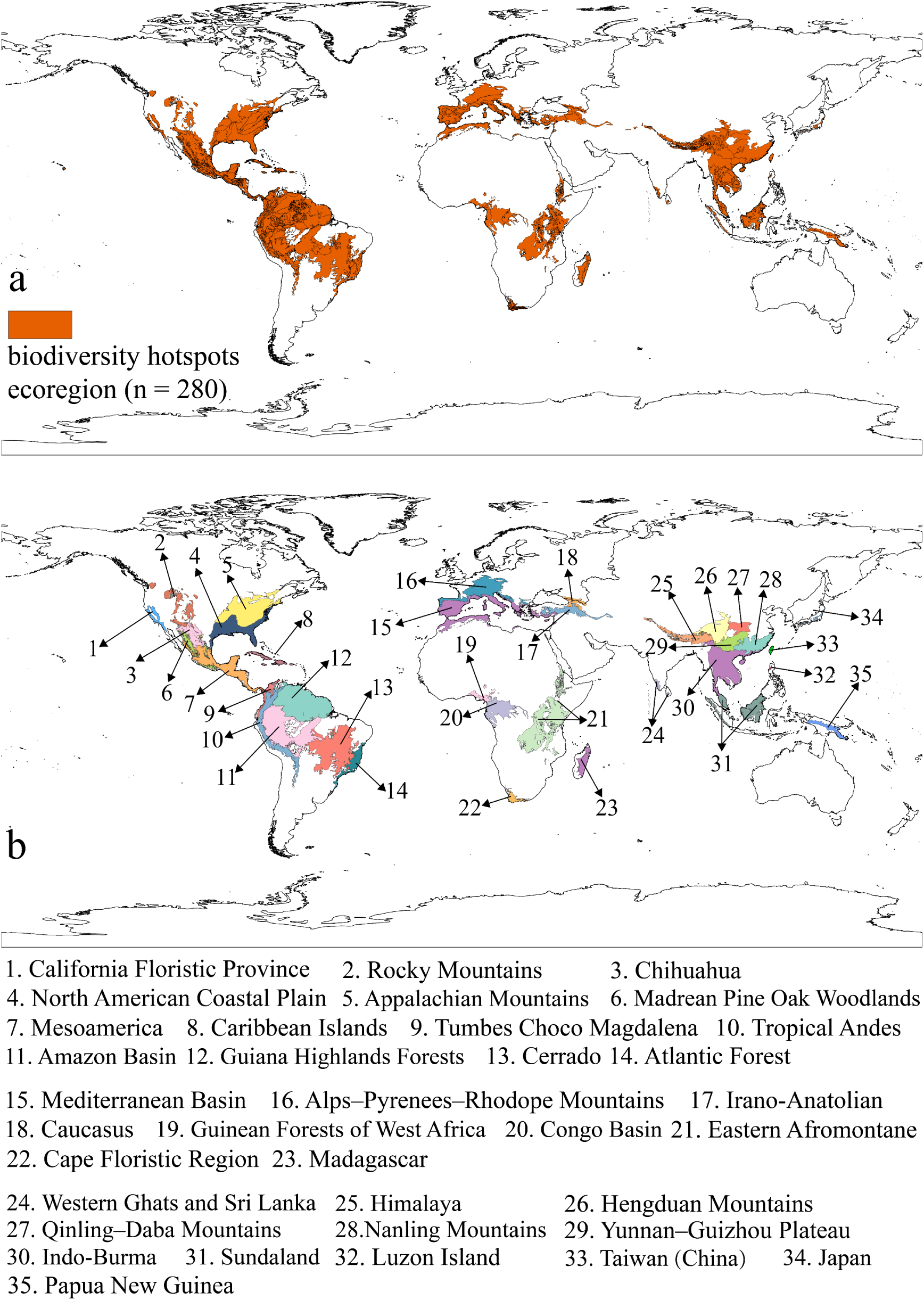
35 global biodiversity hotspots defined by RVP. **a.** The 280 biodiversity hotspot ecoregions included within the 35 global biodiversity hotspots. b. The 35 identified global biodiversity hotspots. The red color highlights 280 hotspot ecoregions, while the other colors highlight different biodiversity hotspots.

### Assessment of Biodiversity Hotspot Identification Criteria

We quantified SPD3 and SPD5 RVP richness for each biodiversity hotspot and assessed whether a minimum richness threshold emerged that identified those ecoregions qualifying as biodiversity hotspots. Simultaneously, under a unified global standards framework, we evaluated whether the proportion of habitat loss and the patch density of native habitats served as additional effective criteria for identifying global biodiversity hotspots.

We first quantified the numbers of SPD3 and SPD5 RVP species within each of the 36 biodiversity hotspots delineated under the framework of Myers *et al*. (*1*) We found that the lower quartile of RVP richness provides a quantitative threshold for hotspot identification: an ecoregion must contain at least 910 SPD3 species or at least 1,120 SPD5 species to meet the minimum criterion (Supplementary Table 1). Nine of the 36 previously identified hotspots failed to meet this threshold, most of which were small island or coastal systems, including the Succulent Karoo, the Eastern Australian forests, New Caledonia, and Polynesia–Micronesia. Among the 35 hotspots identified using our RVP-based approach, the lower-quartile richness values were 1,378 SPD3 species and 1,771 SPD5 species. Only four regions fell below both thresholds (910 or 1,120 species and 1,378 or 1,771 species): the Guinean Forests of West Africa, Japan, the Western Ghats–Sri Lanka region, and Luzon Island in the Philippines.

To quantify native habitat loss across the 36 biodiversity hotspots defined by Myers *et al*. (*1*), we evaluated two unified metrics: (1) the proportion of native habitat loss, defined as the aggregate proportion of non-native habitats and areas exhibiting significant human modification (HM ≥ 0.10); and (2) the patch-density index of native habitats, which measures the extent of landscape fragmentation. We found that six of the 36 hotspots under Myers *et al*.’s framework have lost at least 70% of their native habitat, with the North American Coastal Plain having experienced >80% loss. In contrast, 17 hotspots retained more than half of their native habitat (Fig. 3a). Across these 36 hotspots, patch density of native habitat was positively correlated with the extent of habitat loss, although the two metrics were not identical (Spearman’s r = 0.699, *p* < 0.001; Extended Data Fig. 8), and neither provides a consistent threshold for hotspot identification (Fig. 3a, c). Similarly, among the 35 hotspots identified using RVP, only seven have lost ≥70% of their native habitat, whereas 15 have lost ≤50%. Here too, neither metric provides a universal threshold of habitat loss or degradation for identifying global biodiversity hotspots (Fig. 3; Supplementary Table 1).

**Fig. 3.**
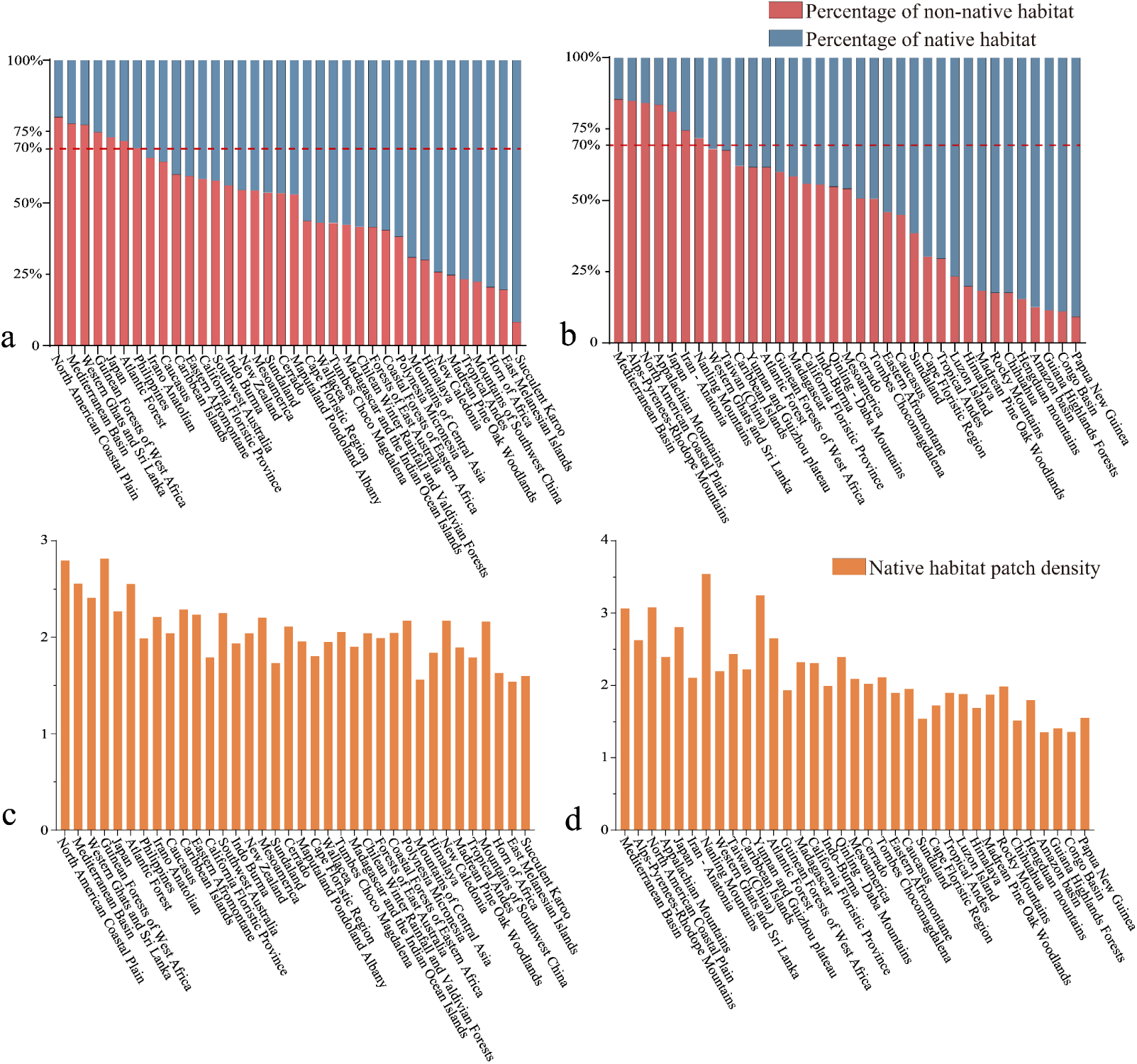
Proportion of native habitat loss based on Myers *et al*.’s Framework and the RVP framework used to determine biodiversity hotspots. (a) proportion of native habitat loss in the hotspots based on the Myers *et al*.’s Framework (36 biodiversity hotspots). (b) proportion of native habitat loss in the hotspots identified based on RVP (35 biodiversity hotspots). (c) patch density of native habitats based on the Myers *et al*.’s Framework (36 biodiversity hotspots). (d) patch density of native habitats based on RVP (35 biodiversity hotspots). RVP = rare vascular plants.

### Global Biodiversity Represented by Biodiversity Hotspots

Due to results based solely on the target of irreplaceability and representativeness of RVP richness (Dobson algorithm) already captured all 36 biodiversity hotspots identified by Myers *et al*., and given the long-standing role of these traditional hotspots in guiding global conservation (*3*, *35*, *36*), we integrated them with the 35 new terrestrial hotspots identified under the RVP framework. During merging, we used the Palearctic–Indomalayan boundary to separate the Japan hotspot (Fig. 2 No. 35) from the Penghu–Taiwan–Ryukyu Islands hotspot (Fig. 4 No. 36). The Papua New Guinea hotspot and the East Melanesian Islands hotspot, share the same biogeographic region and rainforest-dominated ecosystems, and were merged into a single Melanesian Islands hotspot (Fig. 4 No. 43). The integration produced a revised set of 47 global biodiversity hotspots, including 11 new and 17 range-modified hotspots (Fig. 4; Supplementary Table 1). Together, these 47 hotspots cover 26.6% of the Earth’s terrestrial surface, representing a 7.9% increase relative to the 35 RVP-based hotspots and a 9.7% increase relative to the original 36 hotspots under the Myers *et al*.’s framework.

**Fig. 4.**
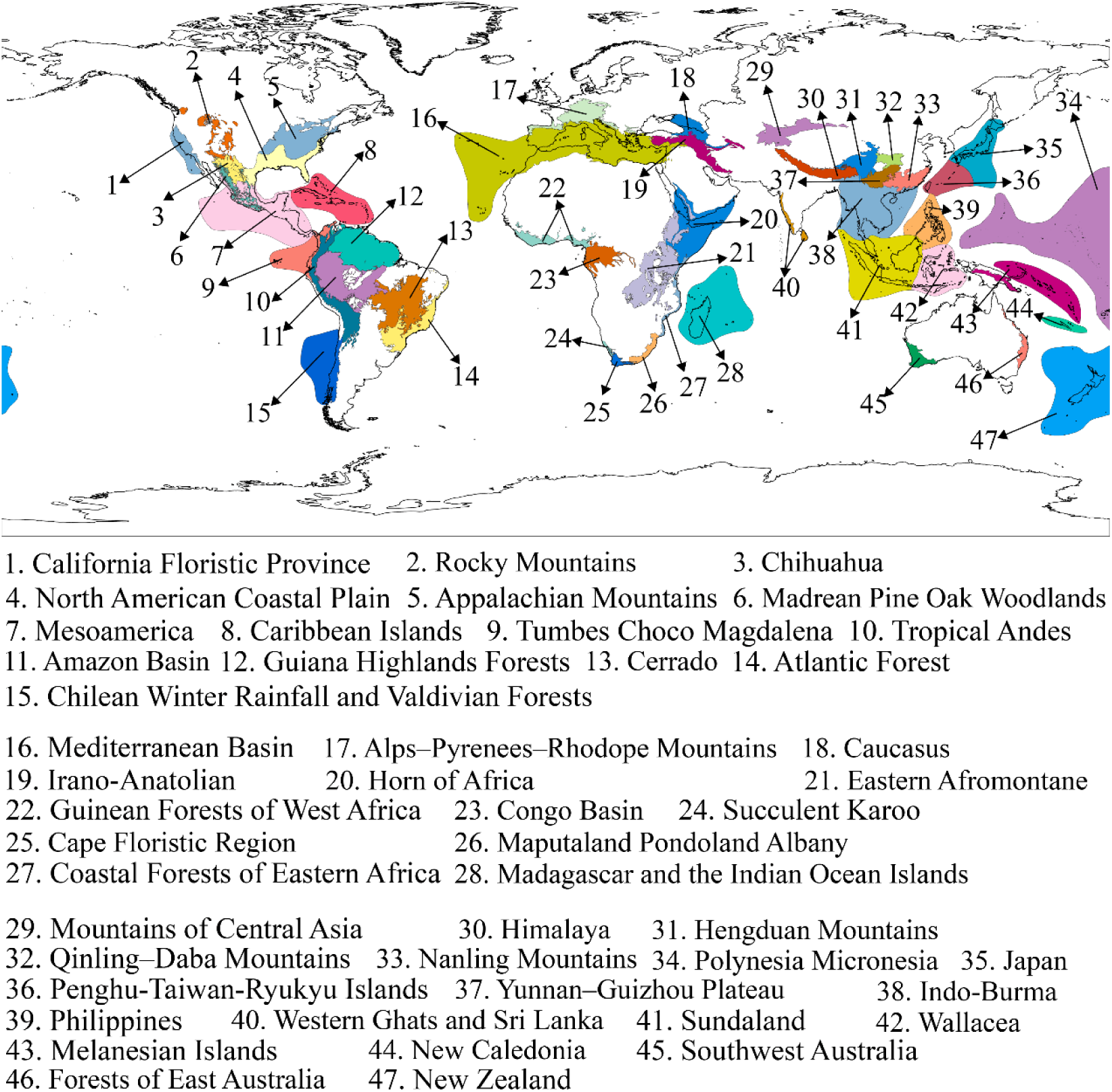
Spatial distribution of 47 global biodiversity hotspots. Colors denote different global biodiversity hotspot regions and their boundaries.

These 47 global biodiversity hotspots encompass more than 80% of RVP, covered under both the SDP3 and SDP5 criteria, and include over 87% of terrestrial vertebrate species of mammals, birds, reptiles, and amphibians (Extended Data Table 1). Notably, the proportion of mammal, bird, and amphibian species within these hotspots each exceed 92% of their global totals. These 47 biodiversity hotspots also contain over 89% of threatened terrestrial vertebrates in the IUCN VU, EN, and CR categories, including 97.4% of threatened birds (Data Table 1, Supplementary Table 3). Among the four terrestrial vertebrate groups (mammals, birds, amphibians, and reptiles) classified as threatened across these 47 biodiversity hotspots, threatened reptiles show the greatest proportional increase, with a coverage gain of 6.2% compared with the 36 hotspots under the Myers *et al*.’s framework (Extended Data Table 1). Moreover, more than 71% of threatened species from the five taxa considered (RVP, mammals, birds, amphibians, and reptiles) have over 90% of their distributional ranges encompassed within these 47 hotspots. Moreover, the 47 biodiversity hotspots redefined based on rare vascular plants (RVP) and three conservation objectives achieve a coverage rate of over 67% for the hotspot areas of threatened terrestrial vertebrates (VU, EN, CR) identified under those same three conservation objectives (Fig. 5).

**Fig. 5.**
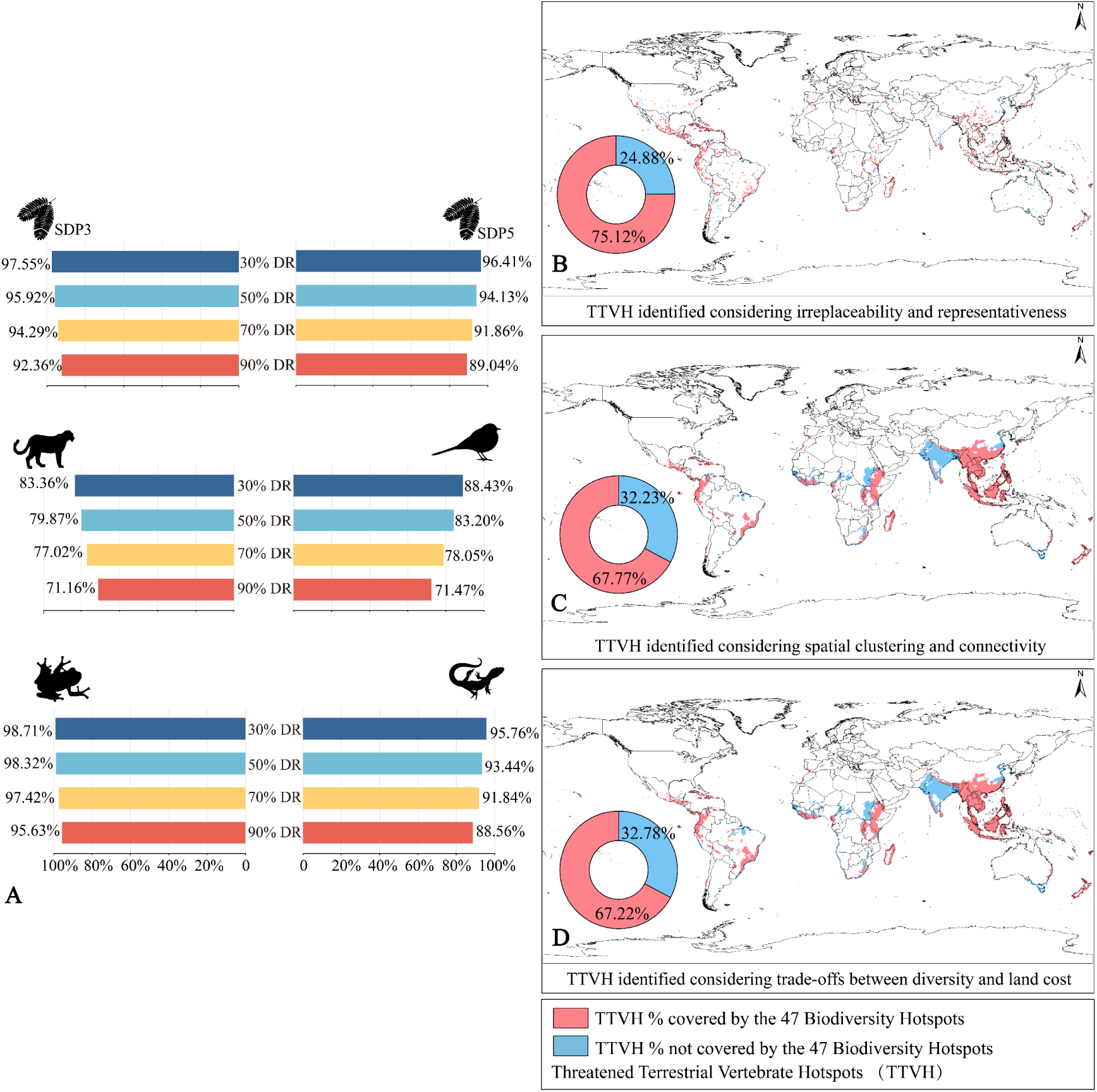
Overlap between 47 biodiversity hotspots (RVP) and threatened terrestrial vertebrate hotspots (TTVHs), and their spatial patterns. (A) The proportion of distribution ranges (DR) for RVP and threatened terrestrial vertebrates covered by hotspots at thresholds of 30%, 50%, 70%, and 90%. (Abbreviations: RVP, Rare Vascular Plants; DR, Distribution Range. Icons: Tiger = Mammals; Bird = Birds; Frog = Amphibians; Lizard = Reptiles; Gymnosperm leaf = RVP). B, C, D Red: Proportion of TTVHs within the 47 biodiversity hotspots; Blue: Proportion of TTVHs within non-biodiversity hotspot regions. (B) Hotspots of threatened terrestrial vertebrates identified by considering irreplaceability and representation. (C) Hotspots of threatened terrestrial vertebrates identified by considering spatial clustering and connectivity. (D) Hotspots of threatened terrestrial vertebrates identified by considering the balance between diversity and land cost.

## Discussion

Over a quarter century of research shows that, owing to shortfalls and biases in global species’ taxonomic and distributional knowledge, no standardized or fine-resolution dataset or identification framework has, prior to this work, reproduced all 36 global biodiversity hotspots defined by Myers *et al.* using expert knowledge (*5*, *9*, *14*). This limitation underscores the shortcomings of previous approaches and the need for more objective, comprehensive, and data-driven methods. By integrating global data on RVP diversity to overcome systematic biases in vascular plant biogeographical information (Extended Data Fig. 6), and by cross-validating results against multiple conservation objectives and identification frameworks, our framework provides a reproducible and globally consistent basis for delineating biodiversity hotspots. We confirm the significance of all 36 previously recognized hotspots, revise the boundaries of 17, and identify 11 additional, previously unrecognized hotspots of critical conservation importance. Collectively, these 47 global biodiversity hotspots encompass 26.6% of Earth’s terrestrial surface and contain 83.8% of RVP, 92.4% of mammals, 96.1% of birds, 87.8% of reptiles, and 95.0% of amphibians. In addition, over 89% of terrestrial vertebrates classified as threatened (VU, EN, or CR) on the IUCN Red List are included. Notably, protecting approximately 27% of the terrestrial land surface could, in principle, safeguard more than 89% of terrestrial species if these hotspots were fully and effectively conserved. Our framework not only reconciles and expands global biodiversity hotspot patterns identified across diverse recent studies using different metrics and taxa (*37–40*), but also establishes a robust, transparent, and scalable foundation for iteratively identifying and refining global conservation priorities as biodiversity data continue to improve. Although spatial patterns of biodiversity can vary among taxonomic groups, our results demonstrate that vascular plant–defined hotspots represent at least 67% of threatened terrestrial vertebrate hotspots. This degree of cross-taxon overlap indicates that plant-based hotspot frameworks retain considerable capacity to reflect broader terrestrial biodiversity patterns. Our findings therefore provide important empirical support for the continued use of biodiversity hotspots as a spatially integrative tool in global conservation planning and investment.

Although our results show that a 910-species threshold for RVP richness is a useful indicator, it should not be regarded as optimal. A global hotspot framework that integrates species richness and irreplaceability with conservation targets will likely provide a more ecologically representative basis for prioritization. By adopting a unified concept of RVP and standardized biodiversity data, our approach reduces reliance on expert-derived estimates of endemic plant richness and distribution, improving both efficiency and replicability (*4*, *41*). At the same time, our revised quantitative standards address long-standing challenges arising from inconsistent criteria for defining endemic plants across regions, as well as from data scarcity. This standardized framework offers a more transparent, repeatable, and ecologically meaningful method for identifying global biodiversity hotspots than what has been used previously.

When evaluating native habitat loss and fragmentation across the 47 biodiversity hotspots, we found that relying on a single numerical indicator as the primary criterion for hotspot identification has substantial limitations. Under the framework of Myers *et al*. (*1*), only six of 36 hotspots have experienced over 70% loss of native habitat, indicating that this threshold is not globally applicable. Cases such as the Madrean Pine–Oak Woodlands and the Rocky Mountains, where habitat loss is relatively low, but fragmentation is severe, highlight the need for multiple assessments of habitat change to better guide policy and management. Relying solely on habitat loss or fragmentation risks overlooking areas that have substantial conservation value due to exceptionally high species diversity (*42*), but have not been significantly disturbed by human activities. Delaying hotspot designation until loss exceeds 70% will dramatically increase restoration costs for ecosystems and threatened species, undermining the goal of preserving more species and intact ecosystems at minimal cost (*43*, *44*).

Significant variation in human impact, remaining habitat quality, and ecosystem diversity across hotspots further suggest that using habitat loss alone will misalign conservation priorities and misallocate resources. For example, the Indo-Burma hotspot has lost 56.1% of native habitat compared with 73.1% in the Western Ghats and Sri Lanka, yet the absolute area lost in Indo-Burma (10%) equals 90% of that lost in the Western Ghats and Sri Lanka. Similarly, assessing hotspots solely by proportional habitat loss neglects regions such as the Amazon Basin, Congo Basin, and New Guinea, where loss is relative less, but ecosystem diversity and integrity are critical for global biodiversity and climate stability. In Brazil, 166,500 km² of tropical forest were lost between 2016 and 2024—an area equivalent to the land area of Paraguay, the entirety of the Western Ghats and Sri Lanka hotspot, or 70% of the Eastern Forests of Australia hotspot (*19*). The Brazilian Amazon is likely approaching a tipping point, which could trigger severe drought and rapid conversion of tropical rainforest to dry forests and savannas (*21*). Once the forest shifts to these alternative states, it becomes highly flammable and can be readily converted to cattle ranches, escalating extinction risk and threatening biodiversity at regional and global scales.

The ambition of Aichi Target 11 to protect 17% of global land by 2020 was only partially realized, leaving one-third of critical biodiversity areas unprotected and less than 8% of land both protected and interconnected (*45*). This shortfall is particularly acute in Asia, where protection reached only 13.2% by 2020, with 73.1% of Key Biodiversity Areas (KBAs) falling short of the 17% target (*46*). Meeting the Kunming–Montreal Global Biodiversity Framework (KMGBF) “30×30” target therefore requires Asian nations to expand protection by 2.4–5.9-fold. However, expanding global biodiversity protection faces several major challenges. First, allocating ∼15% additional land for protection competes directly with agriculture, exacerbating food-security risks in low-income countries that often have rapidly growing human populations (*47*). Second, substantial overlaps between KBAs and Indigenous territories may trigger land conflicts without robust local governance (*48*). Third, conservation financing remains critically insufficient (*49*); for example, African “lion-bearing” reserves face annual funding gaps of USD 0.9–2.1 billion, meeting only 10–20% of operational needs (*50*). Fourth, focusing solely on area targets risks neglecting biodiversity value (*51*, *52*). Between 2018 and 2023, newly established protected areas covered only 7% of threatened species’ habitats, mainly in temperate rather than biodiverse-rich tropical ecosystems (*53*). As of May 2025, seven “biodiversity superpower” countries that support 70% of global biodiversity, including Brazil, Ecuador, and the Democratic Republic of Congo, have yet to submit new National Biodiversity Strategies and Action Plans (NBSAPs) that should have their objectives met by 2030. Of the 36 nations that submitted NBSAPs ahead of COP16, none fully align with the KMGBF 30×30 target (*10*), including Madagascar, Papua New Guinea, and South Africa (*11*). Together, these factors suggest that the feasibility of achieving KMGBF conservation targets and the 2050 “Half-Earth” vision remains limited.

Our results indicate that protecting < 30% of global land area would safeguard > 80% of terrestrial biodiversity, achieving the essence of the 2050 “Half-Earth” vision while requiring substantially less land to do so. The 47 biodiversity hotspots we identified span 184 countries and dependent territories (including 90 high- and middle-income nations), 144 of which have ≥ 30% of their land area classified as hotspots (Supplementary Table 2). Moreover, 9,447 of the 15,507 KBAs (60.9%) located in hotspots collectively encompass 65.4% (6.08 million km²) of the total terrestrial extent of all KBAs (Extended Data Fig. 5). Concentrating conservation action within the 47 biodiversity hotspots offers a disproportionately high return for meeting KMGBF targets. Realizing this potential will require a coordinated set of measures. Directing international and bilateral funding to these hotspots, particularly in the 90 lower- and middle-income hotspot countries, and linking support to NBSAPs with spatially explicit priorities and transparent implementation, would substantially improve protection efficiency. Ensuring Indigenous and local community rights in hotspot regions through the principles of Free, Prior, and Informed Consent (FPIC) and the recognition of Indigenous and Community Conserved Areas (ICCAs) is essential for socially durable conservation outcomes (*54*). Closing the substantial financing gap will require diversified mechanisms, including debt-for-nature swaps, green bonds, payments for ecosystem services, carbon-credit plans, sustainable performance-linked loans (*55*), and blended finance tailored to hotspot nations. Expansion of protected areas should prioritize ecological representativeness and connectivity (*56*), focusing on core zones, ecological corridors, and climate-resilient refugia in hotspots and in corridors to prevent low-quality or poorly targeted growth. Aligning conservation with climate adaptation, sustainable agriculture, and livelihood improvement through cross-sector policy coordination can reduce land-use conflicts and maximize the long-term effectiveness of hotspot-focused conservation strategies.

Although global protected-area coverage continues to expand, prioritizing KBAs within hotspots is essential for delivering meaningful outcomes under KMGBF Target 3, which seeks to conserve 30% of land (*57*). These strategies align with KMGBF principles and instruments, translating the concept of “small land, maximum protection benefits” into actionable international and national policies and establishing a robust foundation for setting achievable Global Biodiversity Framework (GBF) targets for 2050. While such protection is challenging, focusing on safeguarding 80% of global terrestrial biodiversity within hotspots is likely more attainable and scientifically grounded than the “Half-Earth” approach, providing a strong basis for long-term conservation priorities. As biodiversity data and monitoring systems expand, this framework can be dynamically updated to track global conservation priorities in near real time. Quantitative definitions of biodiversity hotspots will be critical for achieving efficient, equitable, and evidence-based protection in the decades ahead.

## Methods

### Data Sources and Database Formation

We compiled a list of 96,140 vascular plant species newly described between 2000 and 2023, based on current taxonomic frameworks for vascular plants—including the Angiosperm Phylogeny Group IV (APG IV) for angiosperms (*58*), the Pteridophyte Phylogeny Group I (PPG I) (*59*) for lycophytes and ferns, and the most recent classification system for gymnosperms (*60*), with scientific names sourced from the IPNI (International Plant Names Index). Using the Global Biodiversity Information Facility (GBIF) application programming interface (rgbif), we systematically retrieved global distribution data for these newly described vascular plants from both specimen and observation sources. Specifically, the specimen data included 16,811,493 distribution records for 46,300 species, while observational data provided 77,679,596 records for 71,133 species. In total, the two data sources contributed 94,491,089 distribution records for vascular plants. The range of each species was delineated using the minimum convex polygon (MCP) method encompassing all occurrence points. In addition, to partition the Earth’s terrestrial surface, we constructed a global hexagonal grid with a side length of 50 km (*61*, *62*). This grid, based on multiple GBIF data sources, was used to estimate the global species richness of newly described vascular plant species (Extended Data Fig. 6). Due to the inherent sampling biases of different GBIF data sources, the species distribution records exhibited spatial skewness and potential spillover effects⁷. For instance, the world’s 20 largest herbaria and natural history museums are concentrated in Europe, the United States, and Japan⁸. In addition, the World Checklist of Vascular Plants (WCVP) project originated in Europe before gradually expanding worldwide⁹. Consequently, the collection and recording of vascular plant species are inherently biased toward economically developed regions with extensive historical surveys.

To address this spatial imbalance and to overcome the challenges of defining and comparing endemic plant species across regions, which are driven by the absence of clear Wallacean delineations and by inconsistencies in scale-dependent endemism assessments (*63*, *64*), we adopted the global plant diversity assessment framework and the concept of RVP proposed by Enquist *et al*. (*27*). Specifically, RVP are defined as those meeting both of the following criteria: (1) ≤3 georeferenced specimen records worldwide (SDP3); and (2) ≤5 records (SDP5). This approach helps to mitigate the effects of an uneven sampling effort and provides a consistent and scalable method for biodiversity assessments across different spatial scales. It also avoids the limitations of traditional endemism-based indicators, which often lack comparability across spatial scales, thereby offering a more objective basis for identifying biodiversity hotspots. We acknowledge that filtering rare species from the raw dataset (vascular plants and their distribution records from 2000 to 2023) may reduce data density and thus weaken the effectiveness of identifying global biodiversity hotspot regions.

The completeness and quality of taxonomic records in the International Plant Names Index (IPNI) have improved significantly, with comprehensive data coverage beginning in 1970 (https://www.ipni.org/about). To enhance data robustness and capture long-term accumulation trends, we expanded the dataset to include vascular plant species newly described between 1970 and 2023. From the IPNI, we retrieved 318,340 species names; after removing duplicates and synonyms, retaining 308,664 valid names published during this period. This represents approximately 88.1% of the total number of described vascular plant species worldwide (*65*). The dataset comprises 292,186 angiosperms, 13,859 ferns and lycophytes, and 2,619 gymnosperms. We then bulk-downloaded global distribution records of these described vascular plant species from 1970 to 2023 via the Global Biodiversity Information Facility (GBIF). We excluded observation records because they are geographically biased (*66–68*), and many consist of AI-identified species rather than specimens collected and taxonomically validated by botanists through rigorous, time-intensive verification (Extended Data Fig. 6a). A total of 20,102,698 georeferenced records were obtained. After removing duplicates and likely erroneous data points, we retained 207,553 distribution records for 125,551 SDP3 species and 345,548 records for 150,487 SDP5 species (Extended Data Table 3).

To evaluate the conservation effectiveness of biodiversity hotspots across multiple taxonomic groups, we incorporated distribution polygon data for four terrestrial vertebrate classes: mammals, avians, amphibians, and reptiles. Distribution data for mammals, amphibians, and reptiles were obtained from the International Union for Conservation of Nature (IUCN; https://www.iucnredlist.org/resources/spatial-data-download), while avian distribution data were sourced from BirdLife International (http://datazone.birdlife.org/species/requestdis). Our vertebrate dataset comprises 5,637 mammal species, 11,175 bird species, 7,949 amphibian species, and 10,047 reptile species.

### Identifying Global Biodiversity Hotspots Based on RVP

To identify biodiversity hotspots, we integrated the spatial distribution of RVP (SDP3 and SDP5) with three distinct conservation objectives and their corresponding analytical methods. Hotspot regions were delineated by integrating the outcomes of these approaches and identifying their areas of spatial overlap. The final hotspot boundaries were refined using a suite of geographic and ecological criteria, including the World Wildlife Fund (WWF) terrestrial ecoregions, the eight major biogeographic realms, recognized geographic subdivisions, and key topographic and geomorphological features (e.g., mountain ranges, plateaus, large river basins, deserts, and island systems) (Supplementary Data Fig. 1, Supplementary Data Fig. 2 and Supplementary Data Fig. 3). In addition, we compiled distribution data for 7,247 terrestrial threatened vertebrates (mammals, birds, reptiles, and amphibians) to quantify global species richness patterns. Threat status followed the International Union for Conservation of Nature (IUCN) Red List categories (VU, EN, CR). Using the same framework applied to rare vascular plants, we delineated terrestrial threatened vertebrate hotspots (TTVHs) under three conservation objectives consistent with those used for biodiversity hotspots based on rare vascular plants, to ensure cross-taxon comparability. We then quantified the spatial distribution and proportional area of TTVHs within the 47 biodiversity hotspots under each objective.

The three conservation objectives and analytical methods are as follows:

1. Considering irreplaceability and representativeness. To ensure complete protection of all RVP in the smallest sites, we applied the Dobson (complementarity) algorithm (*69*, *70*). Specifically, we identified the minimum set of 50-km hexagonal grid cells such that each species was included in only one cell. The process began by calculating species richness for each cell and selecting the cell with the greatest richness. Species within that cell were then marked as covered and excluded from subsequent iterations. The procedure was repeated, each time selecting the cell containing the largest number of yet-uncovered species, until all target species were represented. The resulting set of grid cells was defined as the hotspot region for RVP (Extended Data Fig. 1). This method is inherently a greedy set-cover algorithm and may miss clustered patterns, spatial coherence, and connectivity.
2. Considering spatial clustering and connectivity. We applied Getis-Ord Gi* hotspot clustering analysis to the global species richness of SDP3 and SDP5 RVP (*71*, *72*). Our hexagonal grid system ensures that each spatial unit has two more adjacent neighbors, forming triangular connections that align with the spatial relationships as defined by natural neighborhood methods. Based on this structure, We constructed a spatial weights matrix (SWM) file based on the natural neighborhood contiguity of regular hexagons. Gi* analyses were then performed on the species richness of both SDP3 and SDP5 plants, identifying statistically significant clustering hotspots at confidence levels of 90% and above (Extended Data Fig. 2). This method may be subject to edge effects near study boundaries, and fail to detect fine-scale clusters across broad spatial extents (*73*).
3. Trade-offs between diversity and land cost. To halt global biodiversity loss and species extinctions, Wilson’ s framework advocates protecting at least half of Earth’ s land surface, with the aim of conserving over 80% of terrestrial species (*74*, *75*). This concept underpins the United Nations Convention on Biological Diversity (CBD) 2050 vision (*76*). Building on this framework, we ranked all global grid cells by the richness of RVP (SDP3 and SDP5). For each SDP category, we constructed Pareto curves illustrating the cumulative proportion of species conserved relative to land area protected (*62*, *63*). From these curves, we identified the minimum land area required to conserve at least 80% of RVP in each SDP category and designated these regions as vital biodiversity hotspots (Extended Data Fig. 3). This method relies on empirically chosen thresholds and may result in overlooking spatial patterns (*77*, *78*).

Together, these approaches form a tiered prioritization framework: Dobson ensure irreplaceability, Gi* ensures spatial contiguity, and Pareto ensures conservation efficiency. The integration of these methods captures complementary conservation values, including rarity, clustering, and cost-efficient trade-offs, and addresses the respective shortcomings of the three methods outlined above. Sites identified by both Gi* and Dobson represent high-confidence hotspots. Combining either approach with Pareto solutions provides decision-makers with flexible strategies to achieve the “Half-Earth” conservation target using less land.

To reduce subjectivity in delineating hotspot boundaries, we adopted standardized ecological frameworks. Notably, 24 of the 36 currently recognized global biodiversity hotspots have already delineated their boundaries primarily based on WWF terrestrial ecoregions, or used them as key references for boundary delineation. For example, Conservation International (CI) redefined the boundary of the Mountains of China’s Southwest hotspot to encompass three WWF ecoregions: the Nujiang–Lancang Gorge alpine conifer and mixed forests, the Hengduan Mountains subalpine conifer forests, and the Qionglai–Minshan conifer forests. Similarly, the Eastern Australian Forests hotspot was delineated using the Queensland tropical rainforests and the Eastern Australian temperate forests ecoregions (*41*). To enhance the complementarity and consistency among the three hotspot identification methods, while considering the objectivity of hotspot boundary delineation, we employed a majority rule (≥50) in combination with WWF terrestrial ecoregions to aggregate the six types of RVP hotspot grid cells into hotspot regions with distinct ecological boundaries (Extended Data Fig. 7). Finally, the boundaries of biodiversity hotspots were delineated by integrating ecoregions, topographic features, major biogeographic realms, globally recognized geographic divisions, and climate types. This approach minimizes biases arising from methodological differences in hotspot definition and ensures that each selected region is supported by at least two conservation targets and two analytical methods, thereby enhancing global representativeness and robustness. Although SDP5 RVP numerically encompass those in SDP3, the two categories display distinct spatial patterns with no direct overlap. To preserve spatial heterogeneity resulting from these differences, global biodiversity hotspots were delineated as the union of SDP3- and SDP5-based ecoregion hotspots.

The vector boundary data for the Myers *et al*. framework of 36 global biodiversity hotspots were obtained from the Critical Ecosystem Partnership Fund (CEPF; https://www.cepf.net/). Terrestrial ecoregion boundaries were based on the 867 global terrestrial ecoregions classified by Olson *et al* (*79*). The global climate zonation used in this study was derived from the updated Köppen–Geiger climate classification developed by Peel *et al* (*80*) (Supplementary Data Fig. 2).

### Identification and Assessment of New Biodiversity Hotspots

In 1990, when Myers first proposed 18 biodiversity hotspots(*81*), he defined them as regions containing at least 1,500 endemic vascular plant species and had lost 70% or more of their primary vegetation. We quantified native habitat loss across the revised 36 biodiversity hotspots defined by Myers *et al*. and others using most recent globally consistent datasets on habitat types and human modification. Spatial data on terrestrial habitats were obtained from the UNEP–WCMC *Global Map of Terrestrial Habitat Types* (version 001; ∼100 m resolution; https://data-gis.unep-wcmc.org/portal/home/item.html?id=79134b5187084c2499fc0b1b18e4c6d3), which follows the IUCN habitat classification scheme. Native habitats were defined as all non-artificial vegetation types, including forests, grasslands, scrublands, wetlands, and unused lands, and corresponding to categories 1–8 of the map. Human modification data were derived from the Human Modification Index (version 2; 300 m resolution; https://zenodo.org/records/14502573) (*82*). All datasets were resampled to 100 m resolution and projected to a common coordinate system prior to analysis.

To uniformly quantify the proportion of native habitat loss and the degree of landscape fragmentation, We retained native habitats with no or low human modification (0.00 ≤ Human Modification Index [HM] ≤ 0.10). Accordingly, within each hotspot, non-native habitats and areas with higher human modification (HM > 0.10) were clipped, and the proportion of non-native habitats was calculated. Native habitat fragmentation was quantified as patch density (number of habitat patches per unit area) within native habitat classes for each hotspot using the PyLandStats package (version 2.4.2). To capture local landscape heterogeneity, a 3×3 sliding window was applied to the raster data using rasterio (version 1.3.9). We then used both the proportion of native habitat loss and patch density of native habitats to evaluate the applicability of the threshold of “≥ 70% native habitat loss” as a universal criterion for defining biodiversity hotspots.

After more than two decades of revisions and updates, the number of Myers’ global biodiversity hotspots reached 36 in 2016 (*1*, *4*, *41*, *83*). These hotspots play a pivotal role in guiding the allocation of global biodiversity conservation resources and are widely regarded as credible and valuable in conservation planning. In this study, we used the lower quartile of RVP species richness across the 36 Myers hotspots as the minimum threshold for designating new biodiversity hotspots. This method accounts for the discrepancies arising from divergent definitions of endemic vascular plants across regions, while permitting dynamic adjustments to hotspot thresholds and spatial extents as data availability improves.

During the identification and delineation of new hotspots, we found that most encompassed and overlapped with the 36 biodiversity hotspots identified by Myers *et al*. Consequently, we integrated these established hotspots with our newly identified hotspots to form a consolidated network of 47 global biodiversity hotspots. Finally, to assess how effectively the identified hotspots capture global terrestrial biodiversity, we evaluated the representativeness of the combined hotspot network by quantifying its coverage of the diversity and distribution of major terrestrial vertebrate groups (mammals, birds, amphibians and reptiles), including their threatened species and the distributional ranges of those threatened species (*84*). All calculations were carried out using the R 4.4.2 (*85*) and python 3.8 platform and mapped in ArcGIS 10.8.

## Acknowledgments

This study is dedicated to Norman Myers CMG (1934–2019) and his colleagues for creating the biodiversity hotspots concept and their outstanding contributions to global biodiversity conservation. The year 2025 marks the 25th anniversary of the publication in the journal Nature of “Biodiversity hotspots for conservation priorities (2000)”. We thank Dr. Guopeng Ren for his statistical advice at the early stage of this project. PAG wishes to thank Chrissie, Sara, Jenni, Dax, and Saffron for their support during the writing of this manuscript. This work was supported by the National Natural Science Foundation of China (32471737), the Natural Science Foundation of Yunnan Province (202301AT070247 and 202401BF070001-008), the Candidates of the Young and Middle-Aged Academic Leaders of Yunnan Province (202105AC160070), and the Postgraduate Joint Training Base Project for the Integration of Industry and Education of Yunnan University.

## Author Contributions

Y.Y. conceived the initial idea. Y.Y., RD.W, and X.L. designed the study. X.L. completed data collection and analyzed the data with guidance from Y.Y. and RD.W. Y.Y. and X.L wrote the first draft; R.L. L.B.D., C.A.C. P.A.G. verified the data sources and ensured their accuracy. L.B.D., C.A.C., P.A.G., Y.Y., X.L., G.C.C., RD.W. and R.L. contributed to revising the manuscript, and all authors approved the final version for submission.

## Extended Figures and Tables

**Extended Data Fig. 1.**
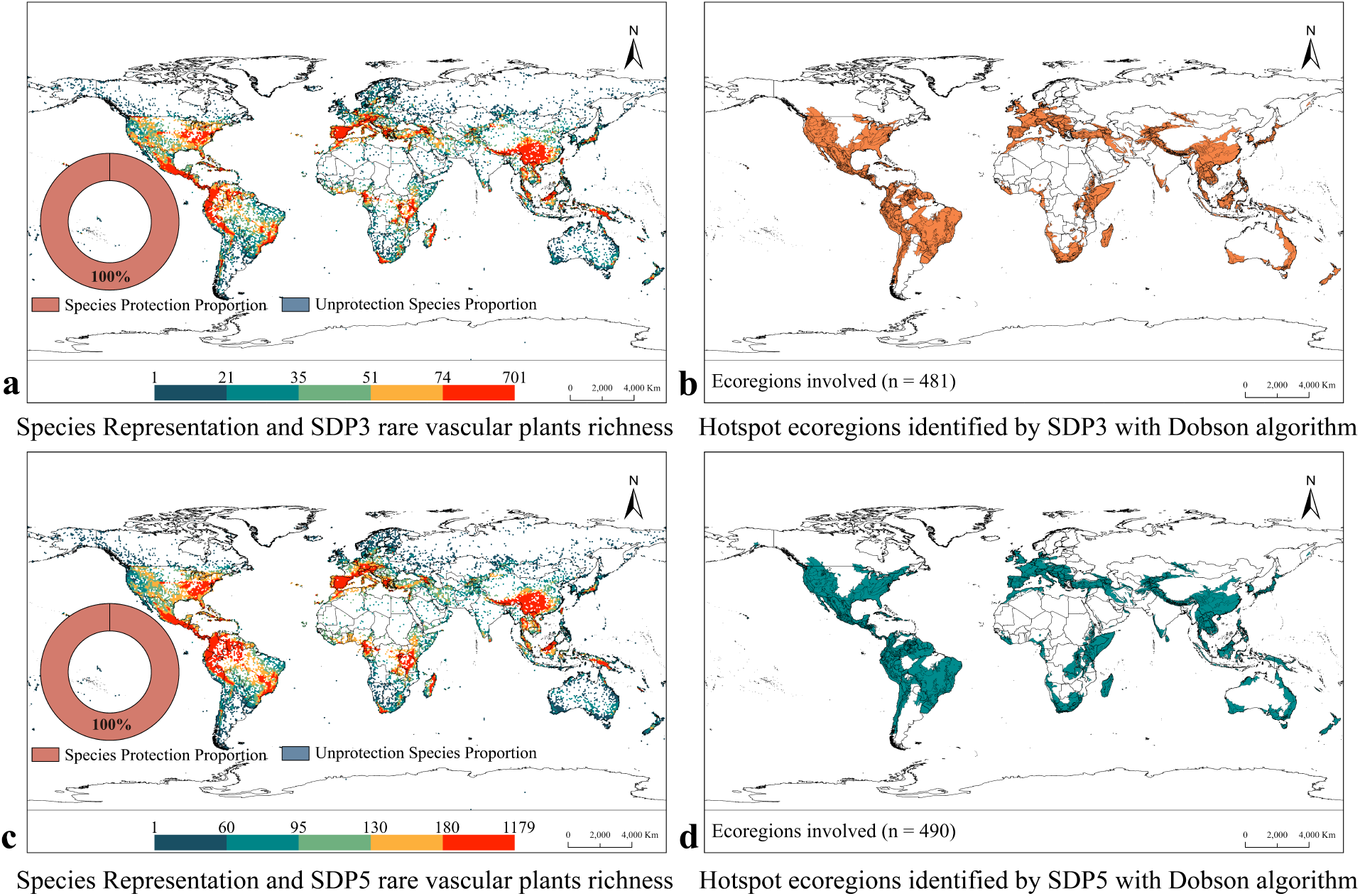
Global hotspots of rare vascular plants (RVP) (SDP3 and SDP5) based on irreplaceability and representativeness. (a) SDP3 hotspots identified under a minimum-area framework ensuring full species representation. (b) Ecoregions involved in SDP3 hotspots. (c) Corresponding SDP5 hotspots identified ensuring full species representation. (d) Ecoregions involved in SDP5 hotspots.

**Extended Data Fig. 2.**
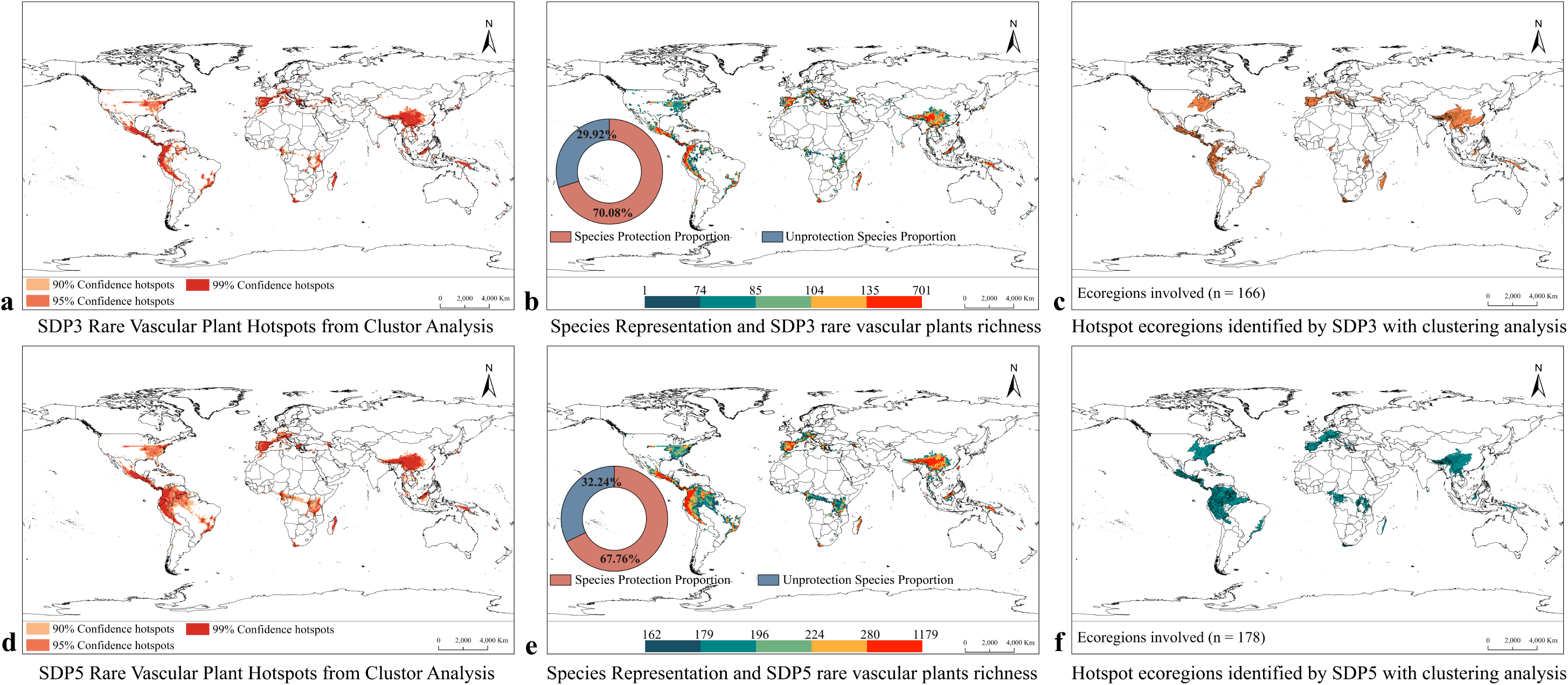
Global hotspots of rare vascular plants (RVP) (SDP3 and SDP5) based on spatial clustering and connectivity. (a), (d) Hotspots for SDP3 (a) and SDP5 (d) identified using Getis–Ord Gi cluster analysis at ≥90% confidence. (b), (e) Spatial distribution of hotspots and corresponding species richness with protection proportion for SDP3 (b) and SDP5 (e). (c), (f) Ecoregions involved in SDP3 hotspots (c) and SDP5 hotspots (f) based on spatial clustering and connectivity.

**Extended Data Fig. 3.**
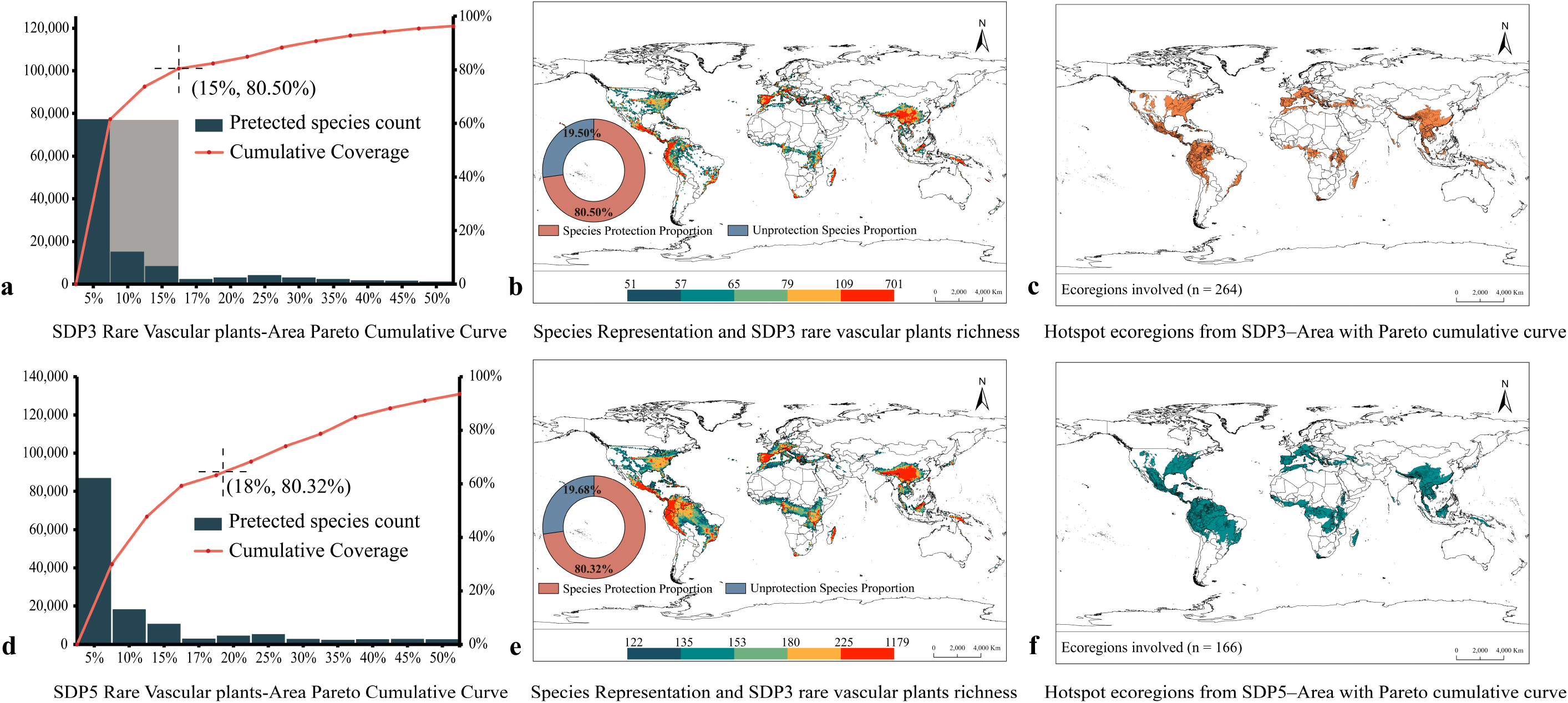
Global hotspots of rare vascular plants (RVP) (SDP3 and SDP5) considering trade-offs between diversity and land cost. (a), (d) Pareto frontier curves showing the cumulative proportion of species conserved relative to land area protected for SDP3 and SDP5, respectively. (b), (e) Spatial distribution of hotspots and corresponding species richness, with protection proportion for SDP3 (b) and SDP5 (e) under the “Half-Earth” target (≥ 80% species protected). (c), (f) Ecoregions involved in SDP3 hotspots (c) and SDP5 hotspots (f) under the “Half-Earth” target (≥ 80% species protected).

**Extended Data Fig. 4.**
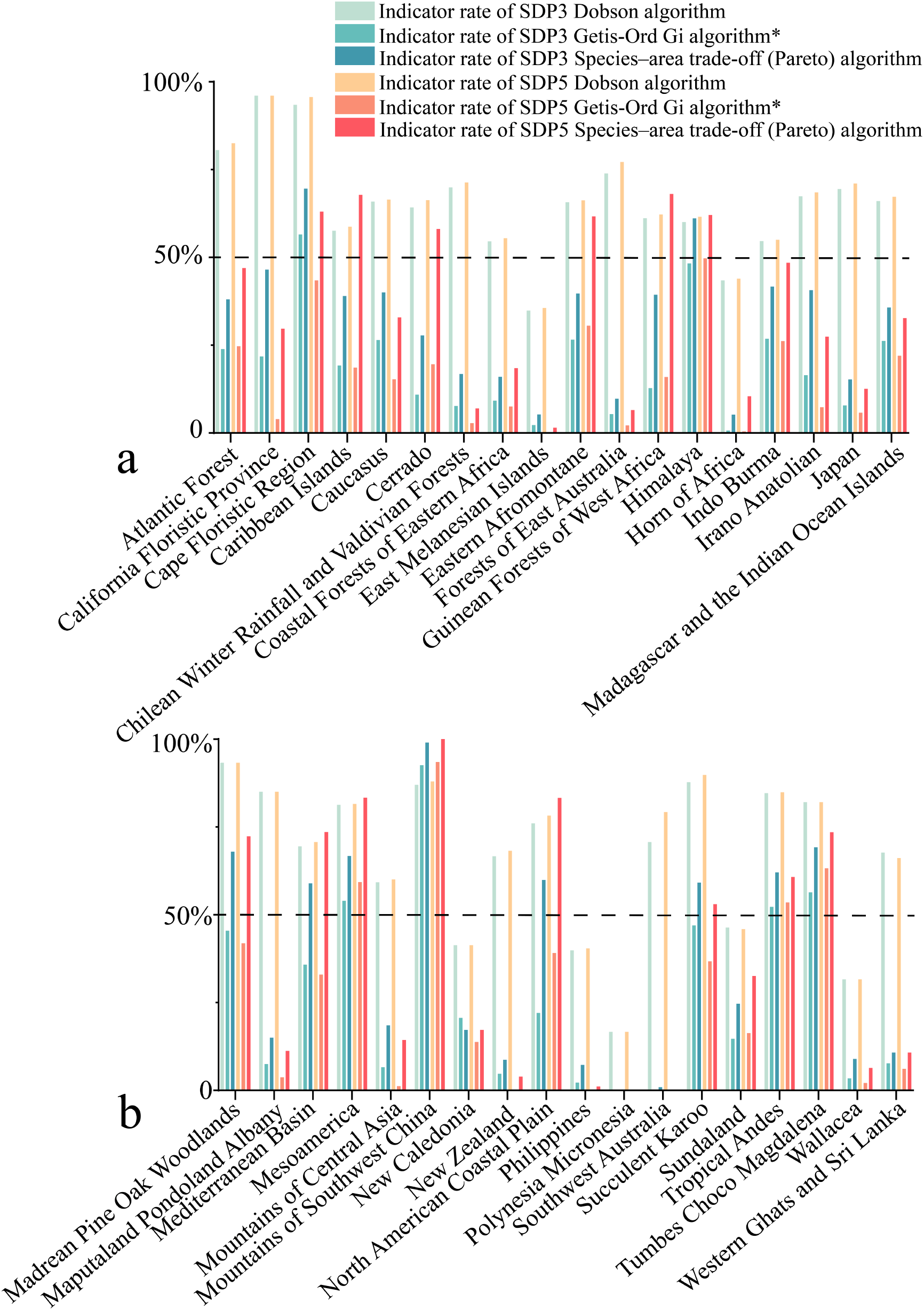
Indicator rates for the 36 biodiversity hotspots under Myers *et al*.’s framework, derived from three hotspot identification methods based on SPD3 and SPD5 rare vascular plants (RVP), expressed as the fraction of hotspot grids identified relative to the total grids defined by Myers *et al*.’s framework.

**Extended Data Fig. 5.**
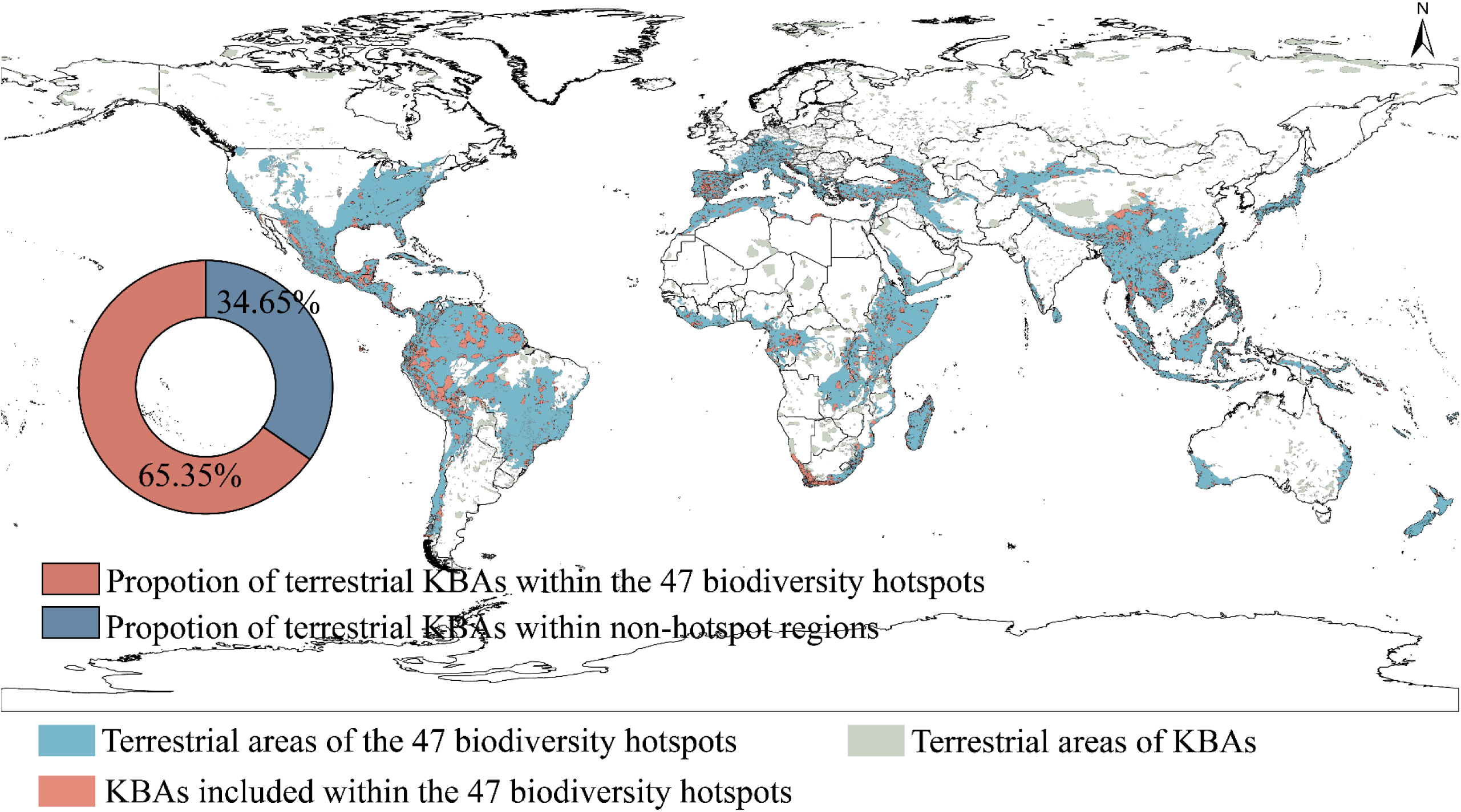
Proportion and spatial distribution of Key Biodiversity Areas across 47 biodiversity hotspots (KBAs). KBA data (2025) accessed from. https://www.keybiodiversityareas.org.

**Extended Data Fig. 6.**
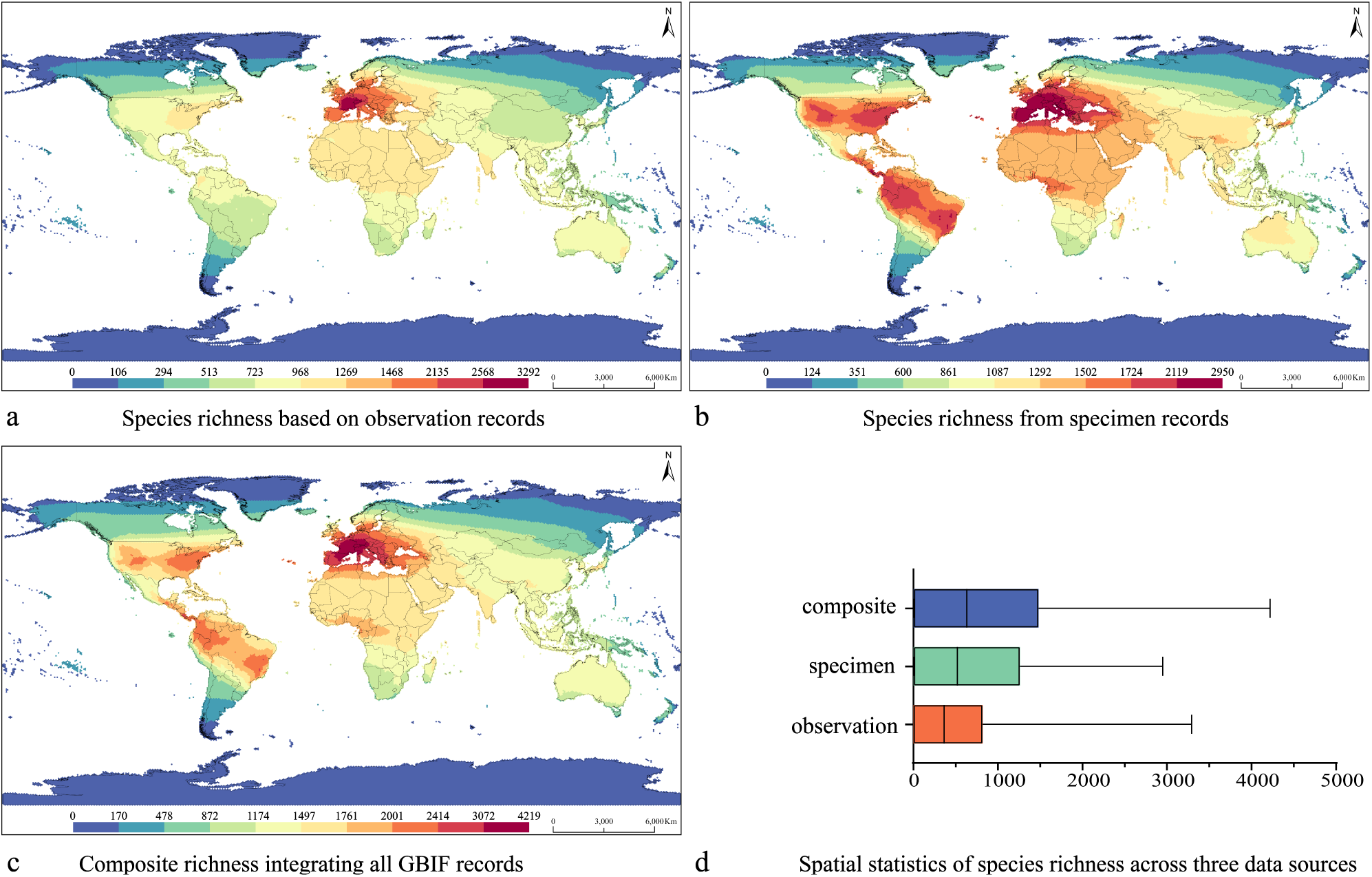
Global vascular plant richness from GBIF (2000–2023). (a) Observation records (field records/AI-derived image recognition). (b) Specimen records. (c) Specimen and observation composite records. (d) spatial statistics: blue=composite; green=specimen; orange=observation. Boxplot representation: center line (median), upper end line whisker (maximum value).

**Extended Data Fig. 7.**
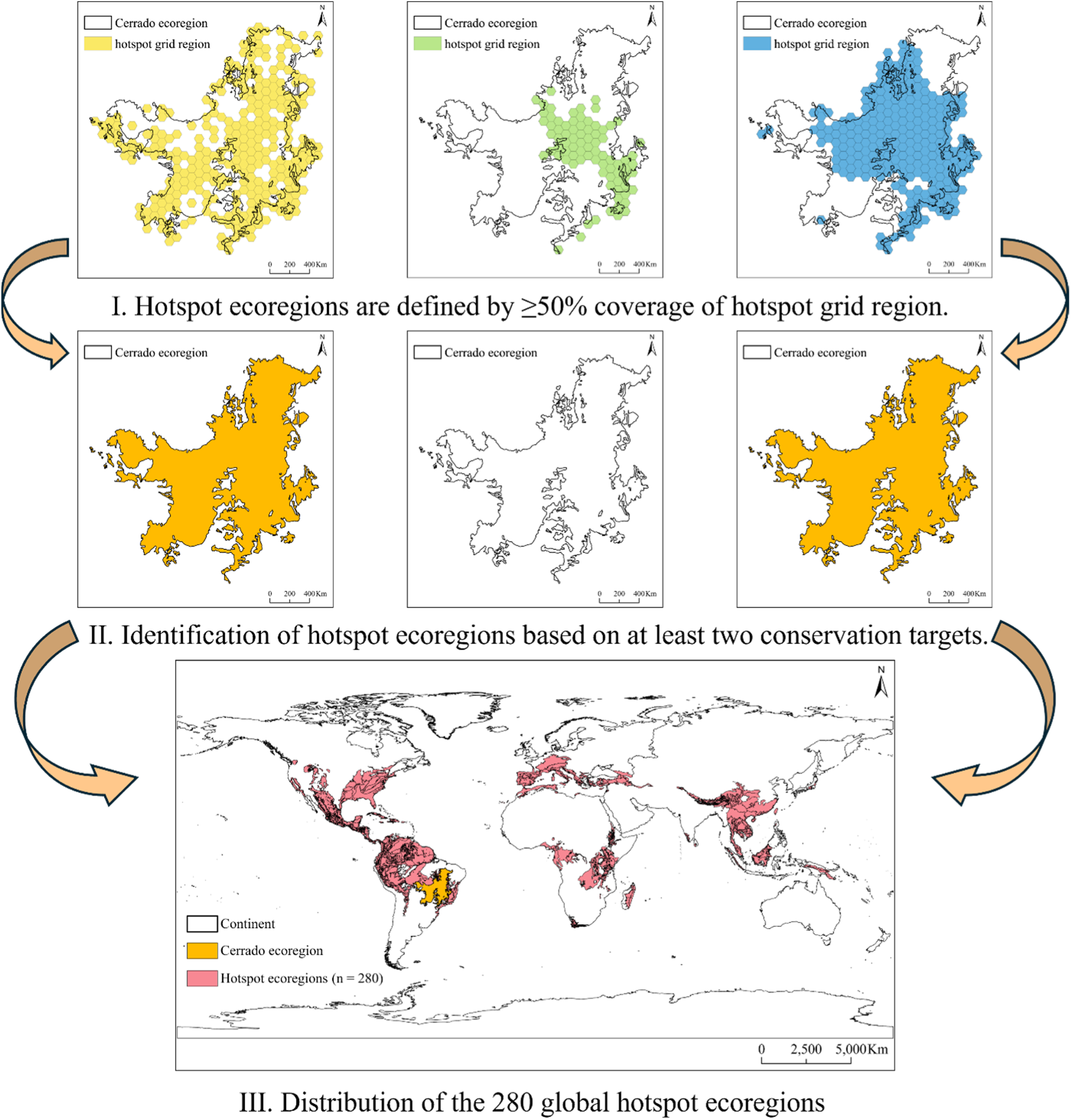
Majority-rule selection of ecoregions in biodiversity hotspot identification based on rare vascular plants (RVP). Hotspots are defined as grid cells with ≥50% overlap with the ecoregion, exemplified here by the Cerrado biodiversity hotspot.

**Extended Data Fig. 8.**
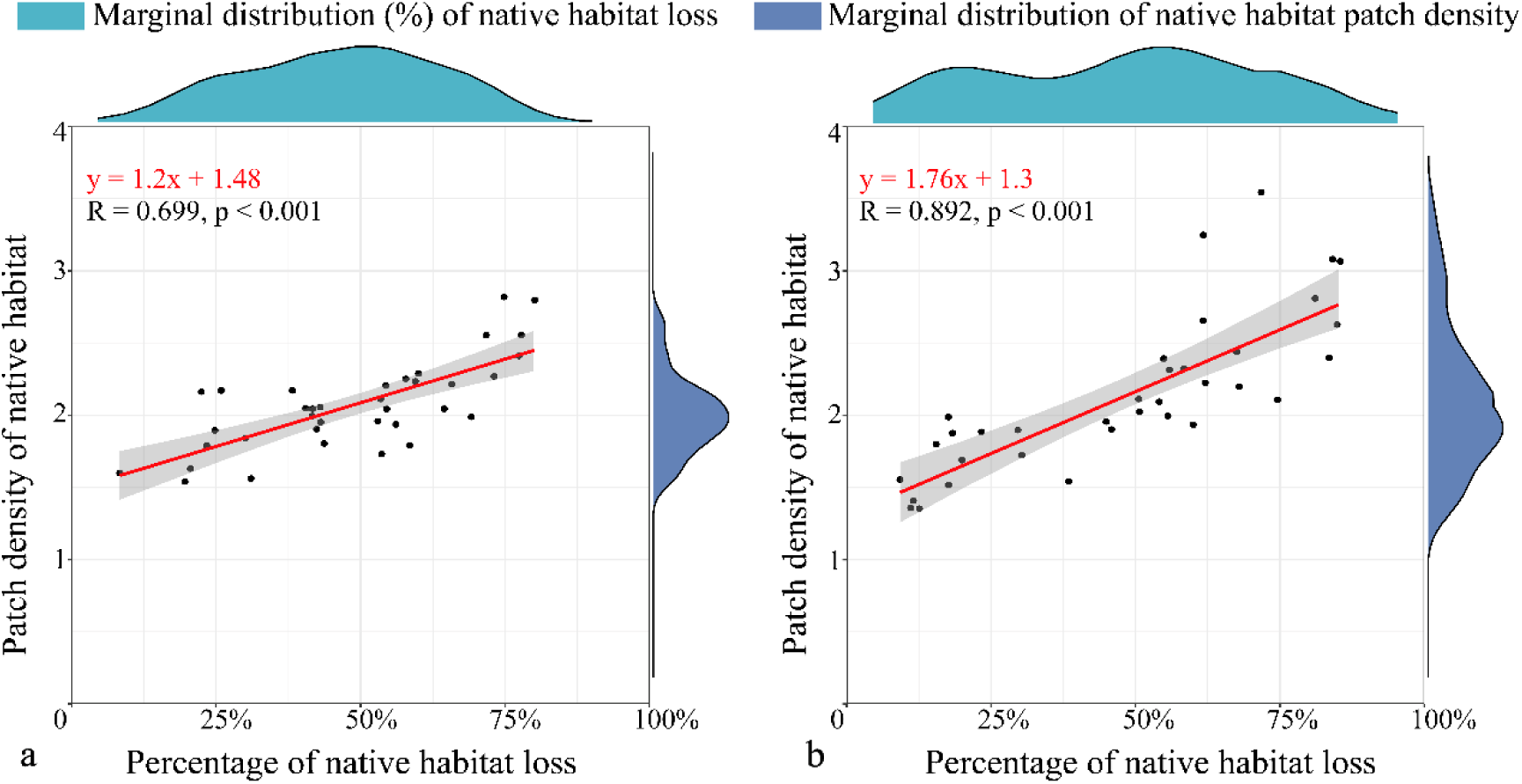
Spearman correlation between the proportion of native habitat loss and native habitat patch density. **a.** Correlation based on the 36 biodiversity hotspots defined under the Myers framework. b. Correlation based on the 35 biodiversity hotspots identified using RVP.

---

**Extended Data Table 1.** Terrestrial area and species coverage across three global biodiversity hotspot categories.

|  | **Terrestrial**<br>**Area**<br>**Percentage** | **Coverage**<br>**Percentage of**<br>**SDP3 / SDP5**<br>**Rare**<br>**Vascular**<br>**Plants** | **Mammal**<br>**Coverage** | **Bird**<br>**Coverage** | **Amphibian**<br>**Coverage** | **Reptile**<br>**Coverage** | **Coverage of**<br>**IUCN**<br>**threatened**<br>**mammal**<br>**species (VU,**<br>**EN, CR)** | **Coverage of**<br>**IUCN**<br>**threatened**<br>**bird species**<br>**(VU, EN, CR)** | **Coverage of**<br>**IUCN**<br>**threatened**<br>**amphibian**<br>**species (VU,**<br>**EN, CR)** | **Coverage of**<br>**IUCN**<br>**threatened**<br>**reptile species**<br>**(VU, EN, CR)** |
| --- | --- | --- | --- | --- | --- | --- | --- | --- | --- | --- |
| 35 hotspots<br>delineated<br>based on rare<br>vascular plants | 18.72% | 66.72% /<br>70.61% | 75.82% | 79.48% | 85.07% | 64.62% | 63.78% | 60.12% | 87.23% | 61.48% |
| 36 hotspots<br>under the<br>framework of<br>Myers *et al.* | 16.97% | 65.56% /<br>67.38% | 84.62% | 90.77% | 83.38% | 82.65% | 84.75% | 88.41% | 91.17% | 86.66% |
| The merged 47<br>biodiversity<br>hotspots | 26.63% | 82.51% /<br>83.78% | 92.32% | 96.08% | 95.07% | 87.58% | 89.83% | 90.93% | 97.38% | 89.85% |

**Extended Data Table 2.** Newly described vascular plant species (2000–2023) from GBIF observation and specimen records, with total species counts after data cleaning.

| GBIF data source | Number of species |  |
| --- | --- | --- |
|  | Raw data | Data cleaning |
| Observation data source | 46,300 | 37,337 |
| Specimen data source | 71,133 | 68,268 |

**Extended Data Table 3.** Distribution records of newly described vascular plant species (1970–2023) from GBIF, and totals after data cleaning.

| GBIF data source | Number of species |  |
| --- | --- | --- |
|  | Raw data | Data cleaning |
| Observation data source | 77,679,596 | 66,564,795 |
| Specimen data source | 16,811,493 | 14,208,596 |
| Total | 94,491,089 | 80,773,391 |

**Extended Data Table 4.** Numbers of vascular plant species and their distribution counts from GIBF specimen data (1970–2023).

| Three major plant divisions |  |  |
| --- | --- | --- |
| Number of angiosperm species | Number of fern species | Number of gymnosperm species |
| 292,186 | 13,859 | 2,619 |
| Total number of species distribution records:16,371,736 |  |  |

**Supplementary Table 1** | Species coverage and native habitat indicators for 36 biodiversity hotspots under the Myers *et al*. (2000) framework, 35 hotspots identified using rare vascular plants (RVP), and 47 integrated biodiversity hotspots, including numbers of RVP, terrestrial vertebrates covered, proportion of native habitat loss, and native habitat patch density.

**Supplementary Table 2** | Income Categories and Area of Countries/Regions within 47 Biodiversity Hotspots

**Supplementary Table 3** | Coverage of IUCN-threatened terrestrial vertebrate species (mammals, birds, amphibians, and reptiles) across three biodiversity hotspot types.

**Supplementary Table 4** | Ecoregions encompassed by 35 biodiversity hotspots identified using rare vascular plants (RVP).

**Supplementary Document 1** | Criteria for identifying 35 biodiversity hotspots based on rare vascular plants (RVP), with associated vegetation and climate types.

**Supplementary Document 2** | Global Biogeographical Regionalization and Climate Types of the 47 Biodiversity Hotspots.

